# Universal transcriptomic signature of age reveals temporal scaling of *Caenorhabditis elegans* aging trajectories

**DOI:** 10.1101/207647

**Authors:** Andrei E. Tarkhov, Ramani Alla, Srinivas Ayyadevara, Mikhail Pyatnitskiy, Leonid I. Menshikov, Robert Shmookler Reis, Peter O. Fedichev

**Affiliations:** Gero LLC, Novokuznetskaya street 24/2, Moscow 119017, Russia; Skolkovo Institute of Science and Technology, Skolkovo Innovation Center, Novaya street 100, Skolkovo 143025, Russia; Central Arkansas Veterans Healthcare System, Research Service, Little Rock, Arkansas, USA; Department of Geriatrics, Reynolds Institute on Aging, University of Arkansas for Medical Sciences, Little Rock, Arkansas, USA; Institute of Biomedical Chemistry, 119121, Moscow, Russia; National Research Center “Kurchatov Institute”, 1, Akademika Kurchatova pl., Moscow 123182, Russia; Bioinformatics Program, University of Arkansas for Medical Sciences, and University of Arkansas at Little Rock, Little Rock, Arkansas, USA; Moscow Institute of Physics and Technology, 141700, Institutskii per. 9, Dolgoprudny, Moscow Region, Russia

## Abstract

We collected 60 age-dependent transcriptomes for *C. elegans* strains including four exceptionally long-lived mutants (mean adult lifespan extended up to 9.4-fold) and three examples of RNAi treatments that increased lifespan by 19 – 35%. Principal Component Analysis (PCA) reveals aging as a transcriptomic drift along a single direction, consistent across the vastly diverse biological conditions and coinciding with the first principal component, a hallmark of the criticality of the underlying gene regulatory network. We, therefore, expected that the organism’s aging state could be characterized by a single number closely related to vitality deficit or biological age. The “aging trajectory”, i.e. the dependence of the biological age on chronological age, is then a universal stochastic function modulated by the network stiffness; a macroscopic parameter reflecting the network topology and associated with the rate of aging. To corroborate this view, we used publicly available datasets to define a transcriptomic biomarker of age and observed that the rescaling of age by lifespan simultaneously brings together aging trajectories of transcription and survival curves. In accordance with the theoretical prediction, the limiting mortality value at the plateau agrees closely with the mortality rate doubling exponent estimated at the cross-over age near the average lifespan. Finally, we used the transcriptomic signature of age to identify possible life-extending drug compounds and successfully tested a handful of the top ranking molecules in *C. elegans* survival assays and achieved up to a +30% extension of mean and median lifespan.

## 1 Introduction

The largest relative lifespan extension yet recorded has been in *C. elegans*, and corresponds to an almost 10-fold increase with the *mg44* nonsense mutation in the *age-1* gene (Ayyadevara, Alla, et al. 2008; Shmookler Reis et al. 2011). However, this hyperlongevity requires homozygosity of the mutation for two generations, resulting in total pre-embryonal genetic disruption. In human subjects, sensible anti-aging therapies would instead be applied in adulthood, ideally at advanced ages. Sadly, the best *C. elegans* models of therapeutic interventions yield significant, but considerably smaller reported increases in life-span by treatments begun at embryonic and post-embryonic stages (e.g., up to roughly +90% by *let-363* RNAi (Vellai et al. 2003)). Late-life pharmacological interventions yielded even smaller effects on lifespan in flies (Moskalev et al. 2010, 2011), nematodes (Ayyadevara, Balasubramaniam, et al. 2017; Ye et al. 2014) and mice (Anisimov et al. 2010; Harrison et al. 2009; Miller et al. 2011). It is not understood why a single nonsense mutation can dramatically extend animal lifespan, while an RNAi block of the same gene does not produce a comparable effect, especially when administered later in life. In many cases, temperature-sensitive mutations extend lifespan without completely eliminating the biosynthesis of the gene product, so the difference is unlikely to be incomplete suppression of transcripts by RNAi. Perhaps the mutation dramatically changes the molecular machinery of the whole organism during development such that the course of aging of the super-long-living strains is qualitatively different both regarding rates and form, and hence could not be easily reproduced therapeutically. Alternatively, perhaps the gene regulatory network is sufficiently robust that a therapy can reduce the pace of aging without qualitative alterations of the relevant molecular mechanisms.

To address these alternatives, we compiled an RNA-seq dataset of age-dependent transcriptomes produced from *C. elegans* isogenic strains and populations that have vastly different lifespans. Among them are three long-lived isogenic strains carrying mutations: *daf-2(e1370), age-1(mg44)* [at the first and second generations of homozygosity], and *daf-2(e1391);daf-12(m20)* double mutant (Ayyadevara, Alla, et al. 2008; Larsen et al. 1995); three RNAi treatments (*daf-4, che-3* and *cyc-1*); and two controls representing wild-type (Bristol-N2, DRM stock). The overall range of lifespans across all the experiments extends from 17 to 160 days. For each of the mutants or interventions, we measured gene-expression levels over time, across their lifespans, collecting 60 transcriptomes in total (9 different biological time-series, each in duplicate).

Principal Component Analysis (PCA) of gene expression reveals aging in all strains and treated groups as a transcriptomic drift in a single direction, consistent across the vastly diverse biological conditions and coinciding with the first principal component of the combined dataset, which is a hallmark of the criticality of the underlying regulatory network (Podolskiy et al. 2015). We therefore expected that the organism’s physiological aging state can be characterized by a single stochastic variable having the meaning of biological age and coinciding approximately with the first principal component score. The aging trajectory, i.e., the dependence of the biological age variable on chronological age, is then universally determined by the underlying regulatory interactions and the experimental conditions through a single phenomenological property describing the effective regulatory-network stiffness. The quantity imposes a natural time scale proportional to the mortality rate doubling time; the fundamental dynamic characteristic of the aging process (Podolskiy et al. 2015).

To evaluate the theoretical model, we performed a metaanalysis of publicly available gene expression measurements in *C. elegans* (more than 4000 samples in total) to identify the aging signature, i.e., the set of genes universally associated with aging across many different biological conditions. We used the same data to introduce a robust transcriptomic biomarker of aging, as a read-out or predictor of “biological age”, and demonstrated its utility across the datasets. The biological age dynamics in our experiments reveal a universal “aging trajectory”: the rescaling of age by lifespan simultaneously brings together the time-dependent trajectories of the transcriptomic biomarker on age and the survival curves. Throughout the paper, “age” means the chronological adult age (post-L4/adult molt for *C. elegans*). Therefore, the universality of aging trajectories may provide a natural molecular basis for the scaling universality of survival curves recently observed (Stroustrup et al. 2016) and independently confirmed in the survival data of all the strains and treatments in our experiments. We investigated the relationship between the stochastic evolution of the biological age variable and mortality using the survival data from an independent experiment. We also experimentally confirmed the model prediction of the equivalence between the mortality rate doubling exponent (inferred at the cross-over age, corresponding to the average lifespan) and the limiting mortality value (corresponding to the mortality plateau). Finally, we used the transcriptomic signature of age to identify possible life-extending drug compounds and successfully tested a handful of them in *C. elegans* survival assays.

## 2 Results

### 2.1 Selection of long-lived strains and life-extending interventions

Several mutations leading to exceptional longevity of *C. elegans* have been identified (Friedman et al. 1988; Gottlieb et al. 1994; Kenyon et al. 1993; Klass 1983) and studied extensively for their remarkable elevations of both lifespan and stress resistance (Ayyadevara, Alla, et al. 2008; Shmookler Reis et al. 2011). We focused on the most long-lived isogenic *C. elegans* strains, carrying mutations in a long-lived wild-type (Bristol-N2 DRM) background: *daf-2(e1370)* [strain SR806], *age-1(mg44)* [SR808, at the first and second generations of homozygosity], and the longest-lived *daf-2(e1391);daf-12(m20)* double mutant [strain DR1694]. The average lifespans in the series range from twofold to nearly tenfold longer than that of the wild type.

To model the effects of therapeutic anti-aging interventions in adult nematodes, we chose five target genes for RNAi inactivation; *daf-4, che-3, cyc-1, cco-1*, and *eat-4.* Mutation or RNAi treatment of each of them had been reported to prolong life-spans: +40-120% by *daf-4(e1364)* (Shaw et al. 2007); +37% by *che-3(e1124)* and +100% by *che-3(p801)* (Apfeld et al. 1999); +87% (Dillin et al. 2002) and +60-100% (S.-J. Lee et al. 2010) by *cyc-1* RNAi; +61% (Dillin et al. 2002) and +57-80% by *cco-1* RNAi (S.-J. Lee et al. 2010); as well as *eat-4* RNAi (Hamilton et al. 2005), for which the percentage increase was not reported.

We confirmed the longevity of worm strains subjected to these five RNAi interventions at 20°C (see Table 1). In some cases, the relative lifespan modification was somewhat smaller, than reported, which was likely due to the different mode of action (mutation or RNAi treatment), or variation in the study protocols, including RNAi-efficacy, exposure times and/or culture temperatures (see Table S1 for comparison). The most substantial relative effect obtained by RNAi corresponds to an increase in median lifespan from 18.5 days to 25 days (+35%) by RNAi of *daf-4.*

**Table 1.** Summary of survivals for the mutant strains and the confirmatory RNAi interventions.

| Mutant strain<sup>a</sup> |  | LS<sup>*</sup>(d) | Rel. LS |
| --- | --- | --- | --- |
| SR806 [*daf-2(e1370)*], N2-DRM background<sup>b</sup> |  | 39 | 2.3 |
| SR808 [*age-1(mg44)*], second-generation homozygotes in N2-DRM background<sup>b</sup> |  | 160 | 9.4 |
| SR808 [*age-1(mg44)*], first-generation homozygotes (“F1”) in N2-DRM background<sup>b</sup> |  | 38 | 2.2 |
| N2-DRM (WT)<sup>b</sup> |  | 17 | 1.0 |
| DR1694 [*daf-2(e1391);daf-12(m20)*]<sup>c</sup> |  | 60.5 | 2.4 |
| N2-DRM (WT)<sup>c</sup> |  | 25.5 | 1.0 |

| Gene for RNAi<sup>d</sup> | Protein encoded | LS<sup>*</sup>(d) | Rel. LS |
| --- | --- | --- | --- |
| N2, FV | (none) | 18.5 | 1.00 |
| ***daf-4* RNAi** | **TGF-beta receptor ortholog** | **25** | **1.35** |
| ***che-3* RNAi** | **Dynein H chain iso 1b** | **24.5** | **1.32** |
| ***cyc-1* RNAi** | **Cytochrome C1** | **22** | **1.19** |
| *cco-1* RNAi | Cytochrome C oxidase | 24 | 1.30 |
| *eat-4* RNAi | BNPI glutamate transporter<sup>**</sup> | 21 | 1.14 |

<sup>a</sup>The long-lived-mutant *C. elegans* strains (Ayyadevara, Alla, et al. 2008; Shmookler Reis et al. 2011) used for mRNA preparation and analysis by RNA-seq. <sup>b,c</sup>The survivals were done separately. The relative lifespan was calculated with respect to the wild-type control from the corresponding series. <sup>d</sup>The survival analysis summary for the RNA interference experiments against the five selected *C. elegans* target genes. Three successful RNAi interventions (in bold) representing apparently different mechanisms of action were selected for further transcriptomic measurements. **Abbreviations:** LS, mean adult lifespan; Rel. LS, relative adult lifespan; N2, the DRM stock of wild-type strain Bristol-N2; FV, empty feeding vector in place of RNAi. \*Mean adult lifespan, days post-L4/adult molt. \*\*Affecting pharyngeal pumping.

### 2.2 Age-dependent transcriptomes of long-lived *C. elegans* strains

The age-dependent RNA-seq experimental dataset consists of the four mutant groups *(daf-2(e1370), age-1(mg44)* [at the first and second generations of homozygosity], and *daf-2(e1391);daf-12(m20)* double mutant), the three examples of life-extending RNAi *(daf-4, che-3* and *cyc-1*, representing knockdown of diverse pathways) and two controls representing *C. elegans* wild-type (Bristol-N2, strain DRM) from Table 1; 60 transcriptomes in total (see Section S4 for RNA-seq data processing details).

We started by performing an exploratory analysis of all the gene-expression data with the help of principal component analysis (PCA). The first principal component (PC1), along which the variance of the data is maximal, is the only component significantly correlated with age (r = 0.75, *p* < 10^−10^, accounting for *r*^2^ = 56% of total variance). PC1 simultaneously arranges the mutants (Fig. 1a) and the RNAi treated strains (Fig. 1b) according to their respective values of chronological age. The total amplitude of change from youngest to oldest ages is approximately the same for all 9 groups despite their wide range of longevities.

**Figure 1.**
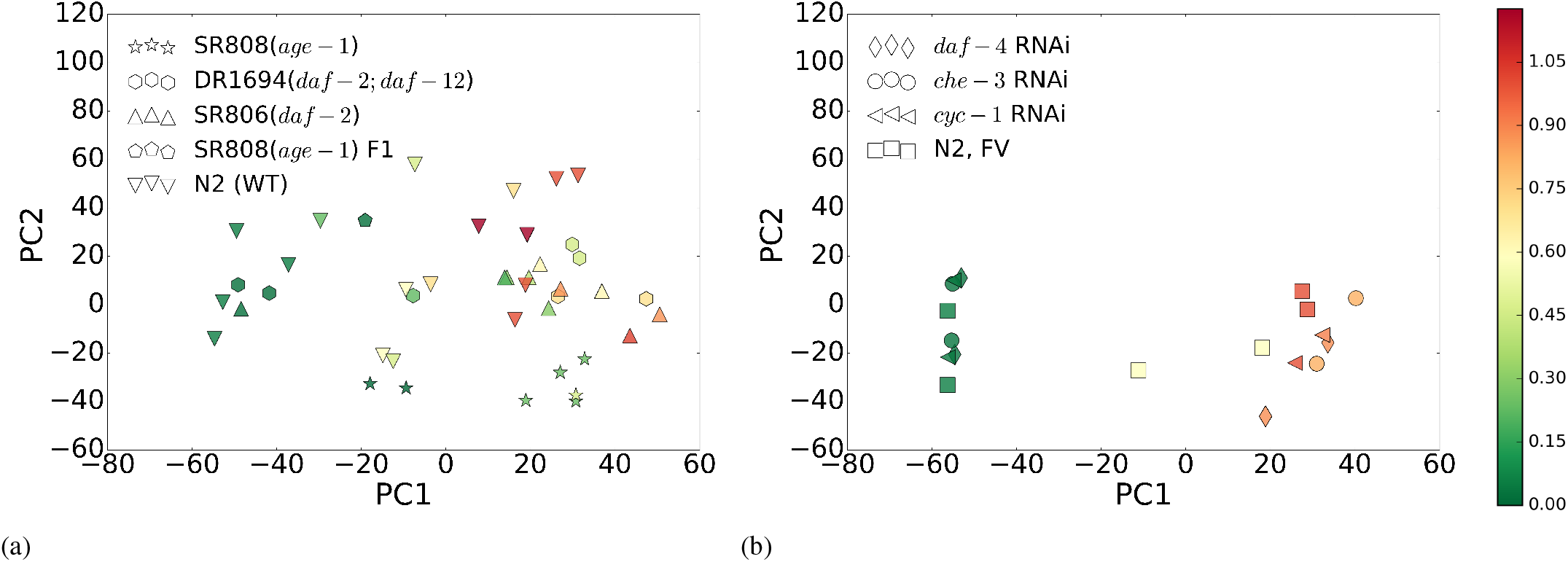
Principal components analysis (PCA) of the experimental RNA-seq datasets for (a) four long-lived mutants and *C. elegans* wild-type (Bristol-N2, strain DRM), and (b) three life-prolonging RNAi-treated groups and *C. elegans* FV controls (fed bacteria harboring empty feeding-vector plasmid). The marker type denotes strains and RNAi groups (see Table 1 for lifespans). The markers in (a) and (b) are colored according to age rescaled to the average lifespan in the groups (see the color bar).

Our gene expression data suggest that the aging signature, i.e., the set of genes transcriptionally associated with age, is robust and remains consistent under a reasonably broad range of experimental conditions and genetic interventions. No other principal component score has a statistically significant correlation with age after correction for multiple comparisons. It is therefore plausible to assume that the state of the organism concerning development and aging can be characterized by a single number, such as PC1 score, indicating normalized biological age. This, however, by no means implies that the system state dynamics along the first principal component alone can explain all transcript-level variation caused by altered biology, mutations or gene silencing. Therefore, the positions of the strain-representing markers in Figs. 1a and 1b can fluctuate along PC1 or in orthogonal directions.

### 2.3 Meta-analysis of aging dynamics in *C. elegans* transcriptomes

Small-scale experiments, including ours, yield a massive number of transcript levels, measured in a relatively small number of samples. The accurate PCA inference of the aging signature and the biomarker of age is challenging due to the lack of consistency of PCA in high dimensions (Gui et al. 2005; Johnstone et al. 2009). The procedure is only appropriate for exploratory purposes since it is prone to overfitting and false-positive errors even if all of the samples are collected in the same laboratory under the same conditions. A direct comparison of gene expression data obtained in different laboratories is further complicated by divergence in experimental procedures, leading to uncontrollable batch effects requiring extensive normalization (Johnson et al. 2007; Lê Cao et al. 2014; Rohart et al. 2017).

To address this hurdle, we proposed that a robust transcriptomic biomarker of age could also be obtained from a sufficiently large collection of publicly deposited “shallow” datasets from small, independent experiments (a dozen samples each, on average) since all the transcriptomes would have to share the same essential biology of aging. In contrast, the specific experimental conditions and uncontrollable batch differences should be mostly uncorrelated, and thus would comprise “noise” in a combined dataset of sufficient overall size.

To investigate such a possibility, we compiled a comprehensive transcriptomic collection for *C. elegans* by combining almost all publicly-available gene-expression experiments into a single database. The resulting “MetaWorm” dataset contains, in total, more than 400 different transcriptomic experiments with *N* = 3724 nematode samples characterized by *G* = 4861 of the most commonly expressed/detected genes in the samples (see Section S5 for more detail on the composition of the dataset and its normalization).

The gene-expression variance in the combined MetaWorm dataset is dominated by batch effects, and hence we do not expect PCA to reveal aging in association with the first principal component in an entirely unsupervised way. Instead, we attempted to identify the aging signature by testing many individual genes for differential expression during aging. Our MetaWorm dataset is sufficiently large to generate the cross-validation ensemble of single-gene association tests using exhaustive random resampling. We further reduced the number of candidate genes by thresholding the transcripts based on the frequency of significant associations in the resampling; we estimate that the chosen crossvalidation threshold corresponds to *p* < 10^−6^ uncorrected, or *p* < 0.005 after Bonferroni correction. The final list of genes robustly associated with aging in the MetaWorm study consists of 327 genes (7% of all genes of MetaWorm): 260 up- and 67 down-regulated with age. We suggest using this gene set as the transcriptomic signature of aging. It is noteworthy that approximately 4000 out of 4861 genes never showed a significant association with aging during the resampling (see Electronic Supplementary Materials for the full list of genes implicated in aging in our calculations).

The transcriptomic signature of age may not be exhaustive, and yet by design, it was reproducible across independent experiments and hence should be useful for future *C. elegans* aging studies. In our experimental RNA-seq dataset, for example, 902 genes are significantly associated with age rescaled by lifespan (for the same threshold as for MetaWorm, *p* < 0.005 after Bonferroni correction) out of 4861 genes most commonly detected in the MetaWorm samples. Even though, the selection using Bonferroni correction is conservative, a list of significant gene-associations in our dataset is larger than for MetaWorm. The difference in the lists presumably occurs due to a laboratory-specific systematic effect on gene expression causing overabundant age-associations, which might not have relation to aging. The prominence of transcription factors among the genes that are age-dependent would inevitably lead to a gene-set with high internal cross-correlation, and a far higher than expected fraction of age-associated genes. The Mann-Whitney U test shows that 327 MetaWorm candidate genes are significantly enriched with the genes having the most significant correlation with age rescaled by lifespan among 902 most significant ones in our RNA-seq data. The corresponding area-under-curve (AUC) statistic for the receiver operating characteristic (ROC) curve is AUC = 0.610 ± 0.015, at the significance level *p* < 10^−30^. This implies that the MetaWorm set of “aging signature” genes very likely includes the same genes that determined PC1 in our RNAseq data, among many more covarying (and hence partially redundant) genes with age-dependent expression.

A correlation with age does not necessarily imply a causal relation to aging, yet genes correlated with age are usually the primary target in aging studies. As a first approach to inference of the regulators of aging, we checked whether the transcriptomic signature of aging is enriched for the targets of known gene-expression regulators (see Table 2). We used four databases for the enrichment analyses: a curated database for transcription factors and RNA-binding proteins from published high-throughput expression studies in *C. elegans*, WormExp (Yang et al. 2016); a high-quality protein-DNA interaction network (Fuxman Bass et al. 2016); and two databases of miRNA-target interactions: the *in silico* predicted TargetScan (Jan et al. 2011) and the experimentally validated MirTarBase (Chou et al. 2016). Enrichment analysis of the list detected ten hits already experimentally characterised as regulators of aging: DAF-16 (Henderson et al. 2001), ELT-2 (Mann et al. 2016), ELT-6 (Budovskaya et al. 2008), PMK-1 (Troemel et al. 2006; Youngman et al. 2011), PQM-1 (Tepper et al. 2014), NHR-1, NHR-10, NHR-86 (Seah et al. 2016), *let-7* (Hsin et al. 1999; Inukai et al. 2013), and miR-60 (Kato et al. 2016).

**Table 2.** Transcription factors, RNA-binding proteins and miRNAs whose targets are significantly enriched in the list of potential transcriptomic biomarkers of age (327 genes).

| Transcription factors or RNA-binding proteins (WormExp)<sup>a</sup> | FDR |
| --- | --- |
| **DAF-16** targets (Kumar et al. 2015) | 4.6e-7 |
| CEC-3 targets (*spr-5* mutant, generation 10) (Greer et al. 2014) | 2.0e-4 |
| small RNAs decreased by starvation at P0 (Rechavi et al. 2014) | 1.3e-3 |
| **PQM-1** L3 targets (Wei Niu et al. 2011) | 1.4e-3 |
| Rb/E2F pathway (DPL-1, EFL-1, LIN-35), intestine (Kudron et al. 2013) | 2.7e-3 |
| **PMK-1** targets down in Day 15 vs. Day 6 (Youngman et al. 2011) | 3.8e-3 |
| age-regulated **ELT-2** targets (Mann et al. 2016) | 3.8e-3 |
| up by CSR-1 and w/out CSR-1-bound 22G RNA (Gerson-Gurwitz et al. 2016) | 7.1e-3 |
| proteins interacting with CEY-1 (Arnold et al. 2014) | 7.1e-3 |
| MUT-14, SMUT-1 (DEAD box) (Phillips et al. 2014) | 1.6e-2 |
| Hox gene targets (LIN-39, MAB-5, EGL-5) (W. Niu et al. 2011) | 3.3e-2 |
| Transcription factors (Fuxman Bass et al. 2016)<sup>a,b</sup> |  |
| **NHR-1, NHR-10, NHR-86** (Seah et al. 2016) |  |
| **ELT-6** (Budovskaya et al. 2008) |  |
| **ZTF-9** (Howe et al. 2016) |  |
| MAB-3 |  |
| miRNA (TargetScan)<sup>a,c</sup> |  |
| miR-57 |  |
| miR-244 |  |
| miR-253 |  |
| **let-7/miR-48/84/241/795** (Hsin et al. 1999; Inukai et al. 2013) |  |
| miR-245 |  |
| miRNA (MirTarBase)<sup>a,d</sup> |  |
| miR-59-3p |  |
| miR-256 |  |
| miR-1819-3p |  |
| **miR-60-3p** (Kato et al. 2016) |  |
| miR-52-5p |  |

<sup>a</sup>Longevity-related regulators are highlighted in bold. <sup>b</sup>The top six transcription-factor hits are reported. <sup>c,d</sup>The top five miRNA hits from each of the databases.

### 2.4 Universal transcriptomic biomarker of age

The robust nature of the gene-expression signature of age across widely varying experimental conditions suggests that at any given time, the organism’s aging state can be characterized by a single number representing biological age. A typical approach for biological age modeling relies on linear regression of measured parameters, gene expression in our case, on chronological age. Naturally, due to the high internal cross-correlation among the gene expression levels and a limited number of samples, the multivariate regression problem is ill-defined, and any number of convenient biomarkers of age could be obtained by applying additional constraints to the regression problem. In this work we impose a requirement of sparsity on the transcriptomic biomarker of age, i.e., the number of genes used in the biomarker model should be minimized while preserving its predictive power. This is possible to do by performing a cross-validated lasso regression of gene expression in the MetaWorm dataset on age rescaled by lifespan in the transcriptomic signature of aging comprising 327 genes. To ensure that the obtained transcriptomic biological age model is not over-fitted and hence retains its predictive power, we have not used our experimental RNA-seq data during training. The final version of the sparse transcriptomic biological age predictor comprises the contributions of only 71 genes (see Electronic Supplementary Materials for the list of regression weights).

### 2.5 Simultaneous temporal scaling of survival curves and aging trajectories

The biological age predictor can now be used to transform our multi-dimensional experimental RNA-seq data representing every sample as a single number, the biological age. In Figs. 2a and 2b we plotted the aging trajectories (the dependence of the biological age on chronological age) and the survival plots. We choose to plot the age-dependent quantities not as a function of age, but as a function of age rescaled by lifespan, in contrast to Fig. S1a and Fig. S1b where the same data are plotted as a function of age without rescaling. The scaling transformation works exceptionally well and simultaneously brings together the survival curves and the aging trajectories of animals with drastically different average adult lifespans: from 17 days for wild-type N2-DRM control worms to 160 days for *age-1(mg44)* mutants (the Pearson correlation of the biological age with age rescaled by lifespan is *r* = 0.86 (*p* = 2 · 10^−18^), cf. *r* = 0.54 (*p* = 8 · 10^−6^) for the correlation of the biological age with age (see Fig. S1a).

**Figure 2.**
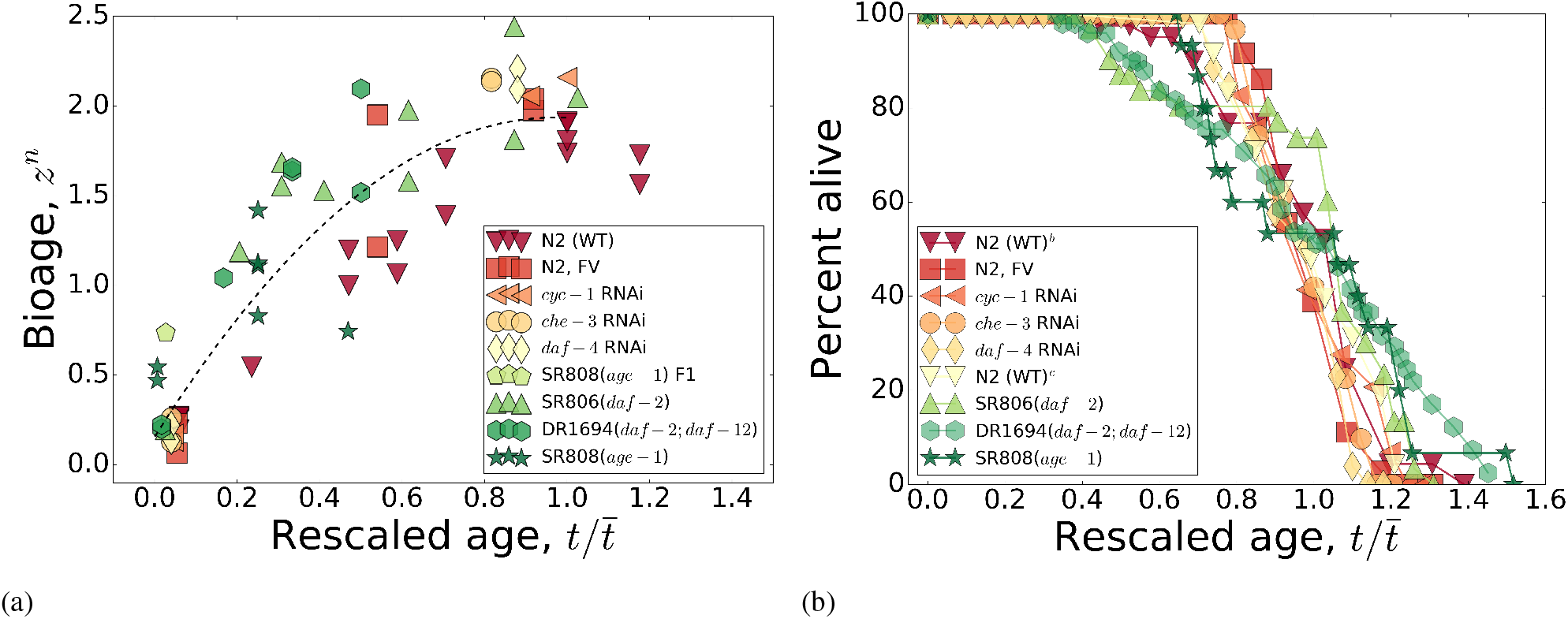
The temporal scaling of survival curves and aging trajectories. (a) The aging trajectories as a function of age rescaled by lifespan for the in-house collection of gene-expression data for four long-lived mutants and *C. elegans* wild-type (Bristol-N2, stock DRM), and three groups of N2 wild-type worms treated with life-prolonging RNAi or bacteria carrying only the empty feeding vector (FV) without an RNAi insert. The dashed line is a guide-to-eye. (b) The survival curves from (a), as a function of age rescaled by lifespan. In both panels, all markers are colored according to the lifespans of the strains (see Table 1 for the lifespans): red for small and green for large lifespans. Overall, the scaling spans almost tenfold variation in median adult lifespan, ranging from 17 to 160 days.

It is worth noting that the biological age naively inferred from the small dataset as a first principal component score from Figs. 1a and 1b is not sufficiently accurate to reveal the temporal scaling of the aging trajectories in the same experiment.

### 2.6 Mortality deceleration

The scaling property of the aging trajectories and the survival functions can be naturally explained using the “aging-at-criticality” model, providing a coarse-grained description of the biological age variable and gene expression dynamics with the help of a simple stochastic Langevin equation and allowing for an analytic solution for mortality and the survival fraction (Podolskiy et al. 2015). Early in life, up to approximately the average lifespan, mortality increases exponentially with age, *M(t)* ≈ *M*_0_ exp *(αt)*, where *M*_0_ is the initial mortality rate (IMR). The “Gompertz” exponent *α* is given by the regulatory network stiffness and is inversely proportional to the mortality rate doubling time (see Section S6 for a summary of the model). We predicted, however, that the exponential rise in mortality rates would cease at late ages, approaching a plateau determined at the value of *α* (Podolskiy et al. 2015).

The mortality records from (Stroustrup et al. 2016) were used to test the theoretical prediction. In that study, nematodes were subjected to various life-shortening stresses and had their lifespans reduced in such a way that any two survival curves could be transformed one into another by a simple rescaling of age. We computed the approximate values of the mortality rate doubling exponent using the data in midlife and the mortality plateau estimates later in life for all the reported conditions (see Section S7 for details of the calculations). The results, summarized in Fig. 3, demonstrate a remarkably tight correlation between the parameters, in good semi-quantitative agreement with the theoretical calculation, across a life-span range of almost two orders of magnitude.

**Figure 3.**
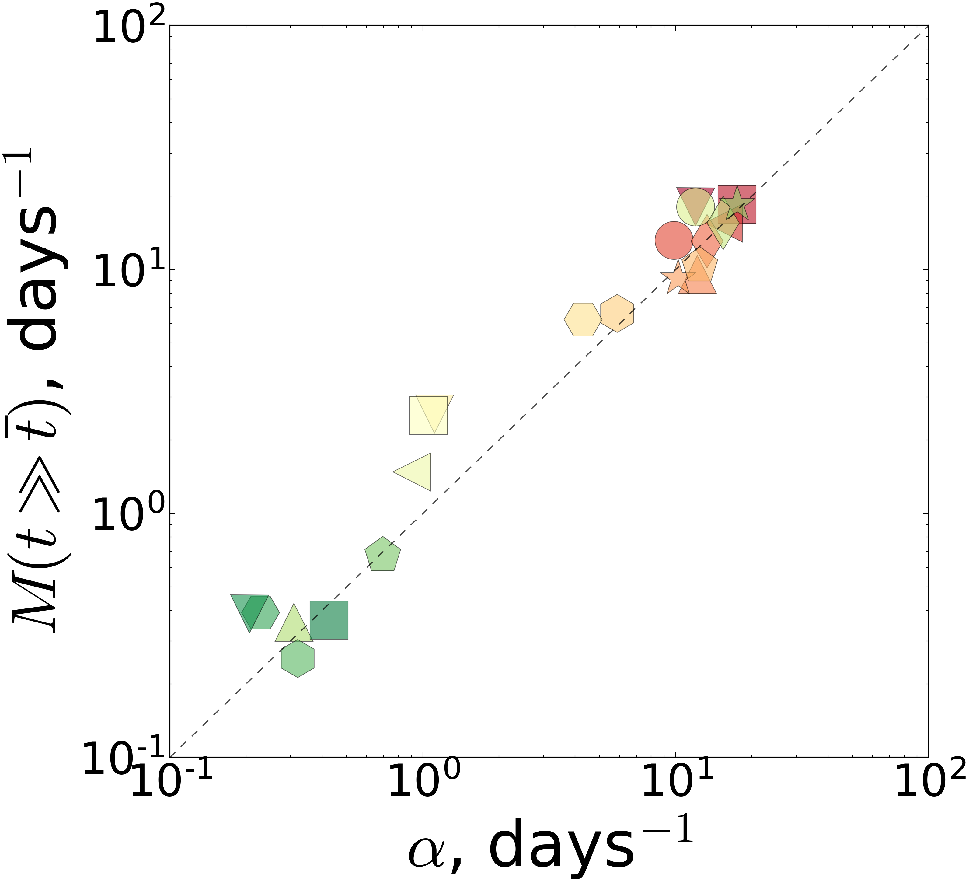
The plateau mortality 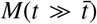 versus the Gompertz exponent *α* calculated from the experimental *C. elegans* survival curves (Stroustrup et al. 2016). The predicted behavior is shown by the dotted line. The values of the average lifespan 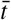 are depicted by the pseudo-color: red for small and green for large values.

### 2.7 Using the signature of aging to identify novel life-extending pharmacological interventions

The results presented so far confirm our association between aging and critical dynamics of the underlying regulatory network. Aging appears to be a consequence of intrinsic instability manifesting itself as lack of dynamic control over the expression of genes comprising the signature of aging. We therefore assumed that interventions exerting perturbations opposing the aging change in the animals would reduce the rate of aging and extend lifespan.

To illustrate the idea, we attempted to identify novel life-extending pharmacological interventions by comparing the signature of aging from this study with gene expression profile changes in response to pharmacologic perturbations from the Connectivity Map (CMAP) database (Lamb 2007; Lamb et al. 2006). This is no trivial task since most of the available transcriptomic studies represent the results of experiments characterizing the effects of drug compounds in human cancer cell strains. We transformed the list of genes associated with aging in *C. elegans* into the form recognizable by CMAP and obtained a list of prospective medicines with gene expression signatures opposing the aging direction and thus presumably capable of reversing the progression of aging in the animals (see Section S8 for the necessary details). A similar approach for drug repurposing against aging has been already demonstrated, e.g., using human brain tissue transcriptomics as the input (Donertas et al. 2018). We observed, however, that the list of the predicted compounds turned out to be sensitive to minor variations in our worm-to-human gene conversion pipeline and the differences between the latest CMAP versions (Subramanian et al. 2017).

From the top drugs in this list, we selected alsterpaullone, metamizole, anisomycin and azacytidine for testing in the *C. elegans* lifespan assay using two concentrations (1 *μ*M and 10 *μ*M) at 20° C; see Table 3 for a summary of the experimental results. All the compounds turned out to be more effective at the lower concentration, which suggests toxicity at the highest dose, probably due to off-target effects. In Fig. 4 we show the survival curves in the respective cohorts. Remarkably, temporal rescaling of survival curves accounts for all variation in survival of the stocks treated with these drugs at both concentrations (compare Fig. 4 to Fig. S2).

**Figure 4.**
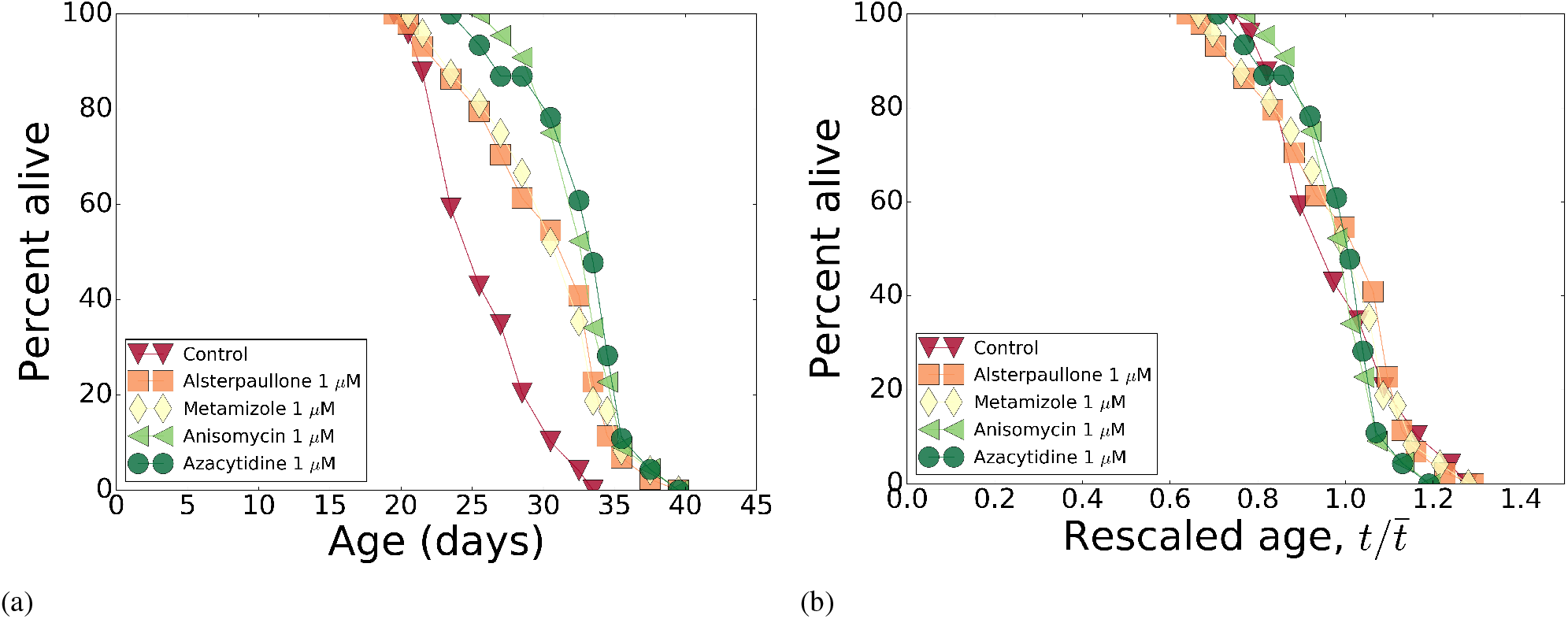
Pharmacological interventions: survivals and their temporal scaling. (a) The survival curves for Bristol-N2 DRM stocks subjected to pharmacological treatments with alsterpaullone, anisomycin, azacytidine, and metamizole at 1 *μ*M; (b) The survival curves from (a), as a function of age rescaled by lifespan. In both panels, all markers are colored according to the measured lifespans of the strains (see Table 3) from red to green for the smaller and for the larger values of lifespans, respectively.

**Table 3.** Summary of survivals for the pharmacological interventions.

| Intervention<sup>a</sup> | Concentration ($\mu\text{M}$) | Mean LS<sup>*</sup>(d) | Median LS<sup>*</sup>(d) | Mean life extension | Median life extension |
| --- | --- | --- | --- | --- | --- |
| Control |  | 26.2 | 25.5 |  |  |
| Alsterpaullone | 1 | 30.6 | 30.5 | +16.6% | +19.6% |
| Anisomycin | 1 | 33.1 | 32.5 | +26.3% | +27.5% |
| Azacytidine | 1 | 33.2 | 33.5 | +26.6% | +31.4% |
| Metamizole | 1 | 30.8 | 30.5 | +17.7% | +19.6% |
| Control<sup>b</sup> |  | 25.0 | 24.5 |  |  |
| Alsterpaullone<sup>b</sup> | 10 | 29.8 | 29.5 | +19.0% | +20.4% |
| Anisomycin<sup>b</sup> | 10 | 25.2 | 24.5 | +0.5% | +0.0% |
| Azacytidine<sup>b</sup> | 10 | 27.7 | 27.5 | +10.5% | +12.2% |
| Metamizole<sup>b</sup> | 10 | 32.5 | 31.5 | +30.0% | +28.6% |
| Control<sup>c</sup> |  | 26.3 | 25.5 |  |  |
| Alsterpaullone<sup>c</sup> | 10 | 28.2 | 27.0 | +7.0% | +5.9% |
| Anisomycin<sup>c</sup> | 10 | 25.0 | 23.5 | -5.5% | -7.8% |
| Azacytidine<sup>c</sup> | 10 | 29.4 | 28.5 | +11.6% | +11.8% |
| Metamizole<sup>c</sup> | 10 | 28.2 | 27.0 | +6.9% | +5.9% |

<sup>a</sup>All survivals were run at 20°C. <sup>b,c</sup>Two independent replicates of the survival experiments. <sup>\*</sup> Adult lifespan, days post-L4/adult molt.
**Abbreviations:** LS, adult lifespan.

## 3 Discussion

The key findings from this study are reduction of gene-expression dynamics to a one-dimension manifold revealed by principal component analysis (PCA), the stability of the aging signature across biological conditions, scaling selfsimilarity of both aging transcript trajectories and survival curves, plateauing of experimental mortality at the predicted level of the Gompertz exponent, and identification of new potential life-extending pharmaceutical treatments.

The observed features of aging dynamics can be explained with the help of an “aging-at-criticality” hypothesis (Podolskiy et al. 2015). This hypothesis proposes that the gene regulatory networks of most species operate near an order-disorder bifurcation point (Balleza et al. 2008) and are intrinsically unstable. In close proximity to the bifurcation, the dynamics of an organism’s physiological state are effectively one-dimensional. Such a reduction of physiological-state vector trajectory in the multidimensional gene-transcript space lets us quantify aging progress by a single stochastic variable representing biological age. This natural indicator of the organism’s aging is directly associated with mortality (see Section S6 for further details). This property of the underlying gene regulatory network is a common feature of complex networks; no matter how complex and large the network is, one can characterize the system by its natural state and control variables, thus effectively describing the system by a one-dimensional nonlinear equation (Barzel et al. 2013; J. Gao et al. 2016). In doing so, one introduces an effective organism-level parameter combining all microscopic features of a network topology into a single number, stiffness or resilience of the network (a counterpart of the rate of aging). The effective state variable is another macroscopic parameter (the order parameter), that plays a role of the aging process indicator and can be ascribed the meaning of biological age.

The apparent stability of the aging signature across the vastly different biological conditions is not surprising from the theoretical perspective since the lifespan and the aging signature are related to the smallest eigenvalue and the corresponding eigenvector of the gene-gene interaction matrix, respectively. Small perturbations, such as effects of mutations or RNAi, thus produce small shifts in the already small eigenvalue and hence may yield very large variations in lifespan. At the same time, alterations of the aging direction by a weak perturbation are expected to be small. The conclusion is rather general and applies to aging in other species, see, e.g., PCA of gene expression levels in normally fed and calorically restricted flies from (Pletcher et al. 2002) and analyzed in (Kogan et al. 2015).

Given the robust and effectively one-dimensional character of changes during aging, a sufficiently large dataset could be used to produce a universal transcriptomic biological age model, such as, in its simplest form, a regression of the gene expression levels on the chronological age, suitable for future aging studies in *C. elegans*. The magnitude and sign of contributions of individual transcripts to the biological age are not unique due to high covariance within each subset of coordinately expressed genes. The model should be fixed by any reasonable additional requirement, such as sparsity, and hence has no direct physical or biological meaning other than providing a convenient tool for experimental data analysis.

The biological age as a function of chronological age defines the aging trajectory and can thus be used to distinguish the progression of aging across strains. Early in life, up to approximately the average lifespan, the age-dependent rise of biological age is a universal function of a dimensionless age variable, obtained by rescaling the chronological age to the strain life expectancy. This is easy to interpret if the influence of stochastic forces is small beyond a certain age, and therefore the progression is almost deterministic with the same time scale defining the shape of the gene expression variations and the value of the average lifespan. In theory, this happens whenever the average lifespan greatly exceeds the mortality rate doubling time. In practice, the assumption is only qualitatively correct, but still provides a reasonable explanation of the experimental situation. According to the model, the temporal scale is defined by the underlying gene regulatory network stiffness and thereby mechanistically relates the organism-level properties, such as lifespan, with potentially modifiable molecular-level interaction properties of the underlying regulatory network, such as the characteristic molecular and genetic damage and repair rates (Kogan et al. 2015).

We expect that the dynamics of gene expression and mortality should increasingly depend on non-linearity of the gradually disintegrating gene regulatory network, as the aging drift and stochastic forces perturb it. Large deviations of gene expression levels from the youthful state are incompatible with survival. Hence the stochastic dynamics of the biological age variable provide a mechanistic coupling to mortality. The scaling universality of the variation in gene transcript levels, along the aging trajectory exemplified by Fig. 2a, should in turn be the molecular basis for the temporal scaling of survival curves, first reported in (Stroustrup et al. 2016). In that study, nematodes were subjected to various life-shortening stresses, and had their lifespans reduced in such a way that any two survival curves could be superimposed by a simple rescaling of age. Our survival data with life-extending mutations, RNA interference, and pharmacological interventions, follow the same pattern. The scaling transformation works exceptionally well and brings together the aging trajectories of animals with drastically different average adult lifespans: from a mere 17 days for wild-type N2-DRM control worms to 160 days for *age-1(mg44)* mutants. Whether the temporal scaling of aging trajectories can be generalized to life-shortening interventions has not been investigated yet, and may be complicated by the multiplicity of pathways whose disruption curtails lifespan.

The temporal scaling of aging trajectories and survival curves is, of course, an approximate statement, since gene expression and lifespans depend on random environmental and endogenous factors. This may be an explanation behind the deviations from the temporal scaling of survival curves in different replicates of the same strain in (Stroustrup et al. 2016). Certain external conditions or therapeutic interventions, in principle may produce a developmental delay or acceleration without a change in the rate of aging, and thus produce deviations from the universal scaling behavior. The DR1694 strain, demonstrating the most discordant survival curve in our analysis, may be an example of such behavior and deserves an independent study.

The biological age should plateau at roughly the average lifespan, which is indeed observed in all our experimental cohorts across a 10-fold range of lifespan difference (Fig. 2a). As the biological age approaches the limiting value, mortality also decelerates and reaches a plateau at the level of the Gompertz exponent obtained from an exponential fit in the age range close to the mean lifespan (Podolskiy et al. 2015). Using the survival data from (Stroustrup et al. 2016), we fully confirmed the theoretical prediction and showed that age-dependent mortality in *C. elegans* indeed decelerates and reaches a plateau at late ages near the expected level. This phenomenon is not limited to the experiments with nematodes and presumably underlies the plateau in mortality rates observed previously in very large populations of medflies and fruit flies (Carey 2011; Rose et al. 2012), which we have extended here to relatively small and homogeneous populations of *C. elegans* (Fig. 3). The results match expectations and, together with the scaling universality of the aging trajectory, both in transcriptomes (Fig. 2a) and corresponding survival curves (Fig. 2b) support our theoretical model.

Multiple explanations have been proposed, for plateauing mortality at advanced ages (Gavrilov et al. 2001; Strehler et al. 1960; Weitz et al. 2001), all involving multi-parametric models. The main advantage of our approach is that, at least in *C. elegans*, the exponential increase of mortality earlier in life and the saturation of mortality late in life are explained within the framework of a simple reaction-kinetics theory dependent only on a single parameter. This parameter is identifiable with the mortality rate doubling exponent measured at midlife on the population level, and with the underlying regulatory network stiffness on the microscopic molecular levels.

Quantification of aging progress using a single number, such as a regression on age of physiological variables representing an organism state, is gaining traction in the aging research community. One of the most successful models of the kind is “DNA methylation age”, which is a weighted sum of DNA methylation features, trained to “predict” chronological age in humans (Hannum et al. 2013; Horvath 2013; Marioni et al. 2015) and mice (Maegawa et al. 2010; Singhal et al. 1987). We note, however, that practical utility of the biological age concept implicitly depends on the assumed lack of variability of the rate of aging in a study. The scaling universality of the aging trajectory reported here suggests that this assumption is not necessarily true, at least in the experiments with *C. elegans*. Nevertheless, the rate of aging is apparently stable in mice (B. G. Hughes et al. 2016) and in humans (Pyrkov et al. 2017), suggesting that lifespan is much more tightly regulated in mammals than in nematodes - consistent with the dearth of spontaneous, very long-lived mutants among “higher” organisms.

The increasing stochastic heterogeneity effects (including the leveling-off of mortality and bioage) help explain when an anti-aging treatment should be applied to obtain the largest possible effect. We speculate that at pre-embryonal and embryonal stages in the simplest animals, or early in life in humans, the growth of an organism is to a large degree determined by a developmental program. At more advanced ages, the stochasticity of the gene regulatory network kinetics takes its toll and leads to increasing phenotypic heterogeneity at every level. Accordingly, we expect that antiaging interventions at the early stages have a broader and more generic effect on aging across diverse species. In contrast, interventions applied at late ages should be precisely selected to help treat specific conditions of an individual at a given point along the aging trajectory; a consequence of life history in the form of stochastically accumulated errors.

The universal aging signature consisting of relatively few genes (less than 7% of all the available transcripts), and these are enriched with the targets of gene-expression regulators that promote longevity via disparate pathways, such as DAF-16, ELT-2, ELT-6, NHR-1, NHR-10, NHR-86, ZTF-9, *let-7*, and miR-60 (see Table 2 for the complete results of the overrepresentation analysis). It would thus be sensible to experimentally test some of the uncharacterized hits from our lists. We tested whether the pharmaceutical interventions (azacytidine, metamizole, alsterpaullone and anisomycin) predicted to exert perturbations opposing the aging change would reduce the rate of aging and extend lifespan, and showed that they indeed prolong lifespan. A version of the extensive LOPAC compound-database with 1280 entries was already screened for lifespan-extending effects in *C.elegans* (Ye et al. 2014). Nevertheless, three of our four hits (metamizole, alsterpaullone and anisomycin) were not tested there, and the fourth (azacytidine) was tested with a negative outcome, which is probably due to toxicity at the higher dose of 33 *μ*M used for the primary screening. In our experiment, all drugs were more effective at 1 *μ*M than at 10 *μ*M, suggesting some toxicity at the higher dose. This is most evident for anisomysin, which is neutral or deleterious at 10 *μ*M.

Alsterpaullone is an ATP-competitive inhibitor of cyclin-dependent kinases (Cdk1/cyclin B, Cdk2/cyclin A, and Cdk5/p25), and with even greater potency, of glycogen synthase kinase GSK-3*β*. Through the latter activity, it inhibits pathogenic phosphorylation of tau in Alzheimer’s disease, and may have other pathogenic targets. Metamizole, or dipyrone, is an inhibitor of cyclooxygenase-3 (Cox-3), observed to activate opioid and cannabinoid receptors; however, it is not considered to be either an opioid or an NSAID. Clinically, it is employed as an analgesic with antipyretic and spasmolytic properties, but only minimal anti-inflammatory effects. It reduces lipopolysaccharide-induced fever (via prostaglandin-dependent and -independent pathways), and disrupts biosynthesis of inositol phosphate. Anisomycin, also known as flagecidin, is a bicyclic derivative of tyrosine that is produced by *Streptomyces griseolus* and inhibits peptidyl transferase activity of eukaryotic ribosomes. It secondarily interferes with DNA synthesis, induces apoptosis in diverse cell types, and is also used as an anti-protozoan agent. It activates stress- and mitogen-activated protein kinases (SAP and MAP kinases) including Jnk and p38/Mapk. Azacitidine is an analog of cytidine, which upon incorporation into DNA (and possibly RNA) irreversibly binds and inactivates DNA methyltransferases. We note that it may inhibit additional targets, e.g. enzymes or transcription factors that bind cytidine or deoxycytidine. Azacitidine and its deoxy derivative, 5-aza-2’-deoxycytidine, are used in the treatment of myelodysplastic syndrome, and of numerous cancers in which anti-oncogenes have been epigenetically silenced. Given the diverse mechanisms of these drugs, they are quite likely to complement one another in a multiple-drug “cocktail”. Moreover, each drug has known, deleterious side effects, which might be avoided or minimized at the low doses evidently required for gero-protection, and especially in drug-combination formulations.

The observed temporal scaling of survival curves and aging trajectories, together with the robust pattern of gene-expression changes associated with aging, appeared to be universal across extremely diverse biological conditions tested in our experiments. From this, we deduce that life-extending effects are achieved by stabilizing the gene regulatory network and by slowing the rate of aging, rather than by qualitatively changing the molecular machinery of the whole organism. This means that the course of aging of the super-long-living strains can be potentially imitated therapeutically, and hence eventually would lead to dramatically increased lifespan without detrimental effects. The “aging at criticality” hypothesis emerges as a robust theoretical and practical framework for the understanding of a broad range of aging-dynamic and survival properties helpful for future efforts to identify anti-aging interventions in *C. elegans* and other species.

## 4 Acknowledgements

We thank G. Getmantsev and O. Burmistrova for proofreading of the work, I. Molodtsov and V. Kogan for invaluable help and advice on data analysis, Prof. Fontana and Nicholas Stroustrup for providing raw survival data and experimental lifetables from (Stroustrup et al. 2016). The work was funded by Gero LLC, NIH Grant P01 AG012411-17A1 (WST Griffin, PI), and VA Merit Review Grant I01 BX001655 (RJ Shmookler Reis, PI).

## Supplementary Materials

### S1 Strains

The following *C. elegans* strains were used in this study: wild-type strain Bristol-N2, subline DRM (herein called “N2” or “N2-DRM”); SR806 *[daf-2(e1370)];* DR1694 *[daf-2(e1391);daf-12(m20)]*, and SR808 *[age-1(mg44)]* at the first (“F1”) and second (“F2”) generations of homozygosity. Strains SR806-SR808 were outcrossed 6 generations into N2-DRM; please see (Ayyadevara et al. 2008) for details. The above mutant strains, and N2-DRM, were grown in 35-mm Petri dishes, on the surface of NGM-agar (1% Bacto-Peptone, 2% agar in nematode growth medium) spotted with E. *coli* OP50 (a uracil-requiring mutant). Several RNAi treatments of genes *(daf-4, che-3, cyc-1, cco-1, eat-4)*, mutation or RNAi of which were reported to prolong life-span, were also assessed. The animals were maintained on NGM-agar plates at 20°C, seeded with *E.coli* HT115 expressing doublestranded RNAs for target-gene knockdowns (Kamath et al. 2003) for both RNA-preparation and lifespan studies.

### S2 Survivals

Lifespan assays were conducted at 20°C, as described previously (Ayyadevara et al. 2008). Briefly, synchronous cultures were initiated by lysis of gravid hermaphrodites in alkaline hypochlorite. Worms were selected at the L4 larval stage, placed 50 worms per plate, and transferred at 1-to 2-day intervals onto fresh plates during days 1-7, and at 2- to 3-day intervals after that. A worm was scored as dead if it failed to move, either spontaneously or in response to a mechanical stimulus; lost worms were excluded (censored) from the survival analysis.

Our survival study confirms longevity of the worm strains subjected to the treatments targeting genes known to affect aging in previous studies. The relative lifespan modification effects in some cases proved to be somewhat smaller, which can probably be attributed to the use of mutation instead of RNAi or a different developmental temperature in the original studies (see Table. S1 for comparison).

Drugs were prepared in small volumes (60 – 100 μl per 10-cm plate), at levels calculated to achieve the indicated concentrations upon equilibration with the full agar-medium volume. Plates were overlaid with drug solutions and rocked with rotation as liquid was absorbed into agar, 24 h prior to use. Worms were transferred to fresh drug-equilibrated plates daily for 12 days and after that, on alternate days (M-W-F).

### S3 RNA isolation

Synchronized strains of *C. elegans* were grown on 100-mm NGM plates, as above, and harvested for RNA extraction at the ages indicated. Worms were washed off plates and rinsed twice in survival buffer; after 30 min at 20°C (to allow digestion of enteral bacteria), they were flash frozen and stored at –80° C. Frozen worms were ground in a dry-ice-cooled mortar and pestle, and total RNA was extracted using RNeasy RNA extraction kits (Qiagen), followed by RNA purification for construction of transcript libraries using TruSeq RNA kits (Illumina, v.2). Sequences are generated as PE100 multiplexes, 100-bp paired-end reads from an Illumina HiSeq2500 or NextSeq instrument, producing 40 – 50 × 10^6^ reads per sample. Paired samples are analyzed with DESeq2 (v1.4.5), and combined sequences are mapped to the *C. elegans* genome using TopHat (Kim et al. 2013).

**Table S1.** Summary of survivals for the mutant strains and the confirmatory RNAi interventions.

| Treatment | Dev. Temp. (°C), | LS, control*(d) | LS, treatment*(d) | Effect on LS | Reference |
| --- | --- | --- | --- | --- | --- |
| *daf-4* RNAi | 20 | 18.5 | 25 | +35% | in-house |
| *daf-4(e1364)* | 20 | 15.8 | 22.1 | +40%<sup>b</sup> | (Shaw et al. 2007) |
| *daf-4(e1364)* | 20 | 14.2 | 31 | +120%<sup>b,**</sup> | (Shaw et al. 2007) |
| *che-3* RNAi | 20 | 18.5 | 24.5 | +32% | in-house |
| *che-3(p801)* | 20 | 18.8 | 37.5 | +100% | (Apfeld et al. 1999) |
| *che-3(e1124)* | 20 | 18.8 | 25.7 | +37% | (Apfeld et al. 1999) |
| *cyc-1* RNAi | 20 | 18.5 | 22 | +19% | in-house |
|  | 25 | 13.6 | 25.4 | +87% | (Dillin et al. 2002) |
|  | 20 | 19.6 | 31.3 | +60%<sup>c</sup> | (Lee et al. 2010) |
|  | 20 | 16.8 | 31.9 | +90%<sup>c</sup> | (Lee et al. 2010) |
| *cco-1* RNAi | 20 | 18.5 | 24 | +30% | in-house |
|  | 25 | 15.2 | 24.5 | +61% | (Dillin et al. 2002) |
|  | 20 | 19.6 | 30.8 | +57%<sup>d</sup> | (Lee et al. 2010) |
|  | 20 | 16.8 | 30.3 | +80%<sup>d</sup> | (Lee et al. 2010) |
| *eat-4* RNAi | 20 | 18.5 | 21 | +14% | in-house |
|  | 25 |  |  | prolonging<sup>***</sup> | (Hamilton et al. 2005) |

<sup>a</sup>The survival experiment for the RNA interference of the five selected *C. elegans* target genes. Three successful RNAi interventions (in bold) were selected for further transcriptomic measurements. <sup>b,c,d</sup>Both experiments were done in parallel. \* Mean adult lifespan, days post-L4/adult molt. \*\* Quote from (Shaw et al. 2007): “*daf-4(e1364)* mutants lived more than twice as long as the wild-type in one trial.”
\*\*\* In (Hamilton et al. 2005), the quantity of effect is not stated, but it is life-prolonging. **Abbreviations:** Dev.Temp., development temperature; LS, mean adult lifespan.

**Figure S1.**
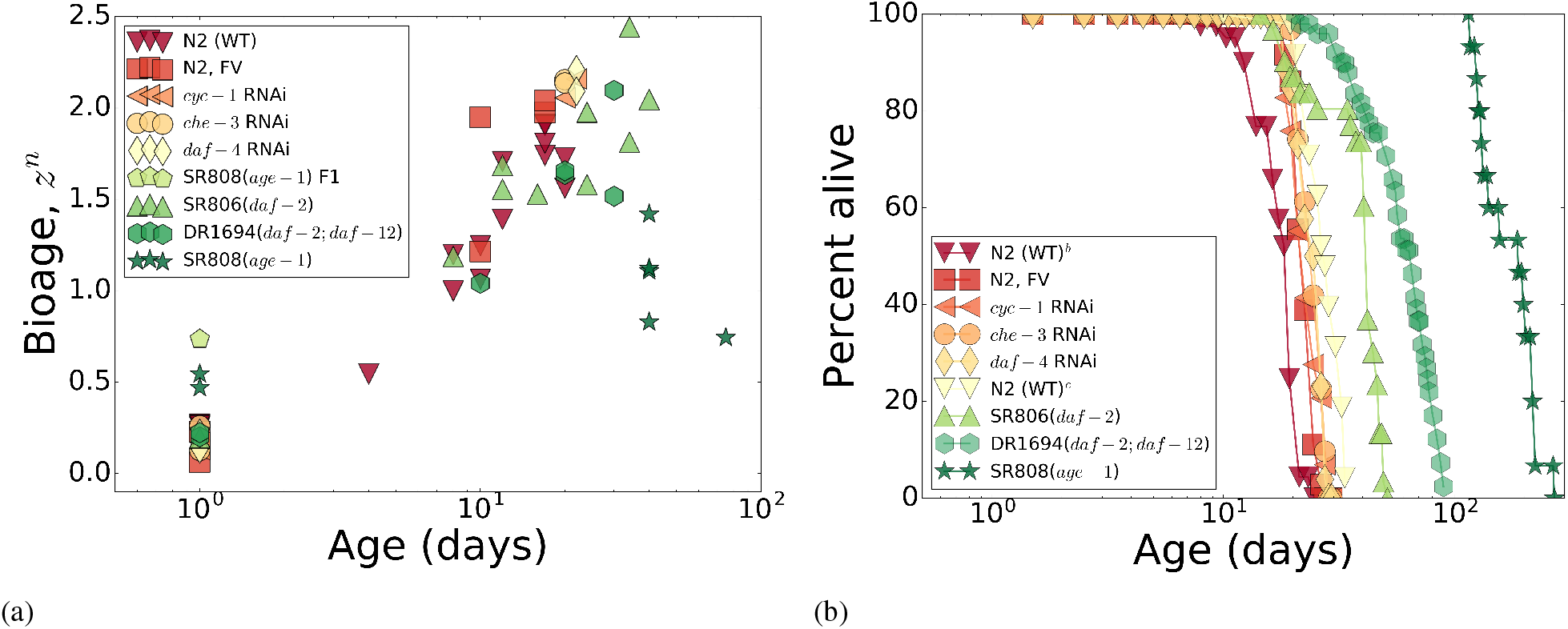
The survival curves and the aging trajectories as a function of age with no rescaling. (a) The aging trajectories as a function of age for the in-house collection of gene-expression data for four long-lived mutants and *C. elegans* wild-type (Bristol-N2, stock DRM), and three groups of N2 wild-type worms treated with life-prolonging RNAi or bacteria carrying only the empty feeding vector (FV) without an RNAi insert. (b) The corresponding survival curves from (a) as a function of age. In both panels, the scale of x-axis is logarithmic, and all markers are colored according to the lifespans of the strains (see Table 1 for the lifespans): red for small and green for large lifespans. Overall, the lifespan ranges from 17 to 160 days.

**Figure S2.**
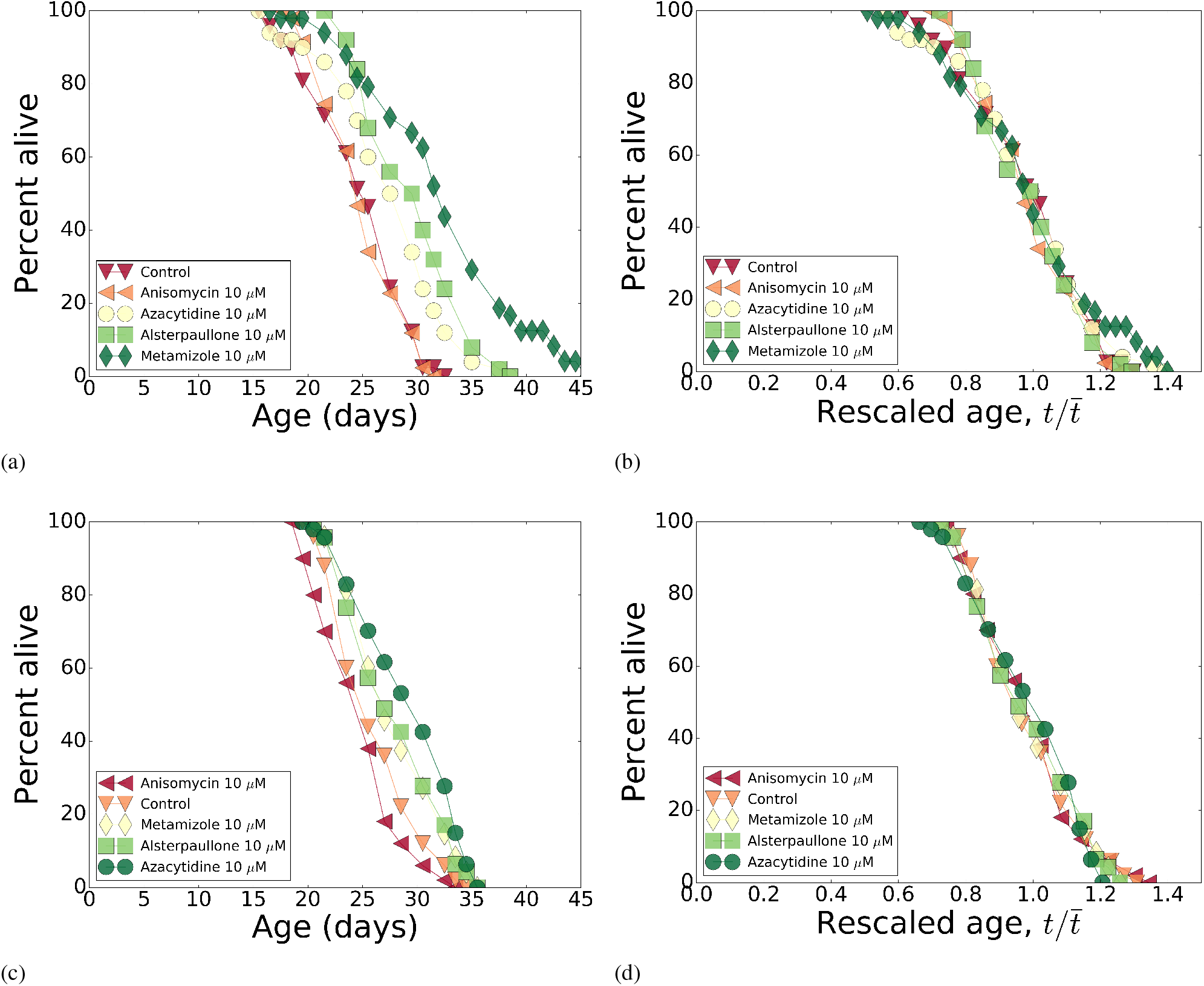
Pharmacological interventions: survivals and their temporal scaling. (a,c) The survival curves for Bristol-N2 DRM stocks subjected to pharmacological treatments with alsterpaullone, anisomycin, azacytidine and metamizole at 10 *μ*M dose. (b,d) The survival curves from (a,c) respectively, as a function of age rescaled by lifespan. In both panels, all markers are colored according to the lifespans of the strains (see Table 3 for the lifespans): red for small and green for large lifespans.

### S4 Experimental RNA-seq dataset

RNA-seq reads were mapped to the *C. elegans* genome (WBcel235, Ensembl annotation) using TopHat 2.1.1 (with --b2-very-sensitive and --GTF options) (Kim et al. 2013) and gene-level read counts were obtained using the htseq-count software (Anders et al. 2015). Low-expressed genes with at least one zero read count per sample were removed from subsequent analysis. Raw read counts were normalized using the upper quartile method (Bullard et al. 2010) and converted to RPKM values using the edgeR library (Robinson et al. 2010).

**Figure S3.**
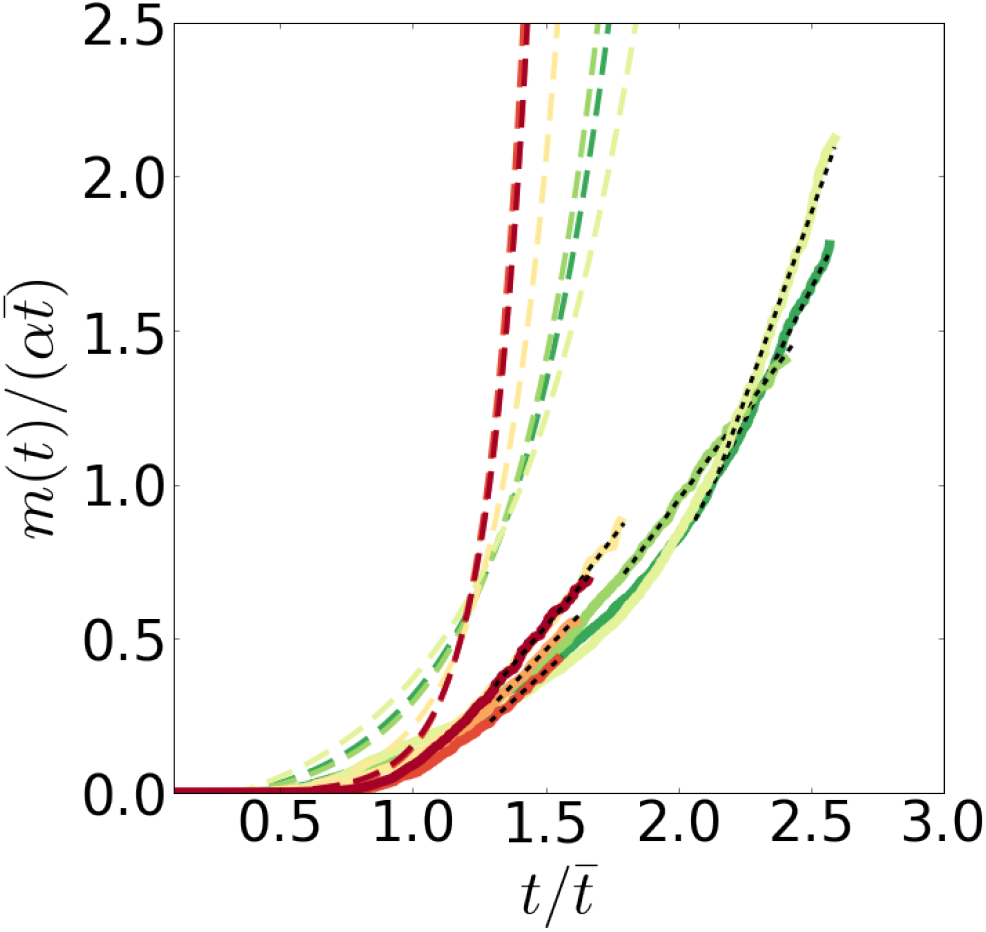
Evidence for the deceleration and plateauing of experimental mortality in *C. elegans*. The normalized cumulative hazard 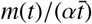 calculated from the Gompertz equation (colored thick dashed) as derived from the experimental Kaplan-Meier plots (Stroustrup et al. 2016) (colored thick solid lines). The tail of the cumulative hazard (black thin dashed lines) was used for the calculation of the plateau mortality 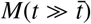 by linear regression. The values of the Gompertz exponent *α* are indicated by the pseudocolor: red for large and green for small values.

### S5 MetaWorm dataset

The “MetaWorm” dataset was compiled from almost all publicly available information on expression profiles *C. elegans* from GEO database (Barrett et al. 2013) and annotated with the corresponding worms’ ages and strain lifespans. For individual genes represented by multiple probesets, the probeset with the largest signal was used. Gene expression in all datasets was normalized using the YuGene (Lê Cao et al. 2014) algorithm, which was specifically developed for gene expression comparisons among different platforms. The final dataset represents a 3724 × 4861 matrix (samples-x-genes) and incorporates more than 400 transcriptomic experiments (see Electronic Supplementary Materials).

### S6 Critical dynamics of gene regulatory networks

We focus on transcriptomic data and describe time-evolution of gene expression by a matrix 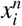, where indices *n* = 1… *N* and *i* = 1… *G* enumerate samples and gene transcripts, respectively, *G* is the total number of genes and *N* is the total number of samples. The measurements are taken at successive instances of time/age, *t^n^*. Following (Podol skiy et al. 2015), we characterize the gene-expression kinetics by a differential equation: *dx_i_*(*t*)/*dt* = *f*(*x_i_, t*), where all the kinetic properties of an organism representing the gene-gene interactions are encapsulated into the function *f* (*x_i_, t*). The coarse-grained description of aging dynamics can be obtained from the linearized version, 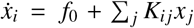, assuming small deviations from the steady state. Here *K_ij_* = *df_i_/dx_j_* is the matrix of interactions among the genes. The stability properties of this matrix determine, whether the corresponding gene regulatory network would be stable (all eigenvalues of *K_ij_* are negative), or unstable (at least one eigenvalue is positive).

In systems without evident symmetries, the system’s dynamics phenomenology near the critical point, separating stable and unstable regimes (critical behaviour), is that of codimension-one bifurcation. More specifically, there is only one of all negative eigenvalues of the interaction matrix *K_ij_* approaching zero and becoming small and positive, *α* > 0. The system’s kinetics are mostly associated with fluctuations along the right eigenvector *b_i_* of the matrix *K_ij_*, corresponding to this eigenvalue. The gene expression variation is dominated by the critical mode associated with the singular eigenvalue of the interaction matrix *K*. Therefore, the transcriptome of aging animals can be accurately described by a one-factor model

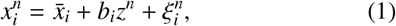

where 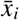 is the initial system state. The data covariance matrix is highly singular, 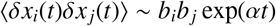 and hence the mode vector *b_i_* coincides with the first principal component, see Figs. 1a and 1b.

The mode variable *z^n^* is the order parameter with the meaning of biological age, which follows the stochastic Langevin equation:

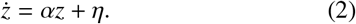

Here the random variable *η* represents the stochastic effects of external and endogenous stress factors. Over time, on average, the solution of the stochastic equation describes exponential deviations from the initial point, subsequent disintegration of the gene regulatory network, and, eventually, death of the organism.

The characteristic time scale in Eq. (2) is defined by the underlying gene regulatory network stiffness *α* and thereby mechanistically relates the organism level properties, such as mortality rate doubling time and lifespan, with the regulatory network topology quantified by potentially modifiable molecular-level interactions variables encoded in *K_ij_* and characterizing molecular and genetic damage and repair rates (Kogan et al. 2015). Small perturbations modify the gene-gene interactions and produce a change of already small eigenvalue, *α*, and hence may result in huge variations in the lifespan. At the same time, alterations of the aging direction, *b_i_* by the very same weak perturbation would remain small. Therefore, we expect that aging trajectories cor responding to different lifespans are self-similar and different by a single time scale factor *α*.

Practically, one calculates biological age by projecting the gene expression data into it with the help of a transcriptomic biomarker of age, *a_i_*

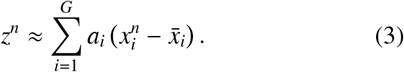

The definition of the transcriptomic biomarker *a_i_* is not unique, since any vector, orthogonal to *b_i_* can be added to the projector *a_i_* without changing the prediction results significantly, especially if the experimental noise (such as batch effects) is large. The best possible candidate for the transcriptomic biomarker of age *a_i_* is the left eigenvector of the interaction matrix *K_ij_* corresponding to the eigenvalue *α*.

### S7 Mortality analysis

The discrepancy between the mortality behavior predicted by the Gompertz equation and experimental mortality for late ages in *C. elegans* is sufficiently large and thus can be used to test the aging theory predictions quantitatively with high-quality mortality data. Mortality data of appropriate quality were recently published in (Stroustrup et al. 2016), where a temporal scaling law of aging in *C. elegans* was observed, similar to that inferred for *D. melanogaster* (Helfand et al. 1995). This scaling law states that under the influence of some intervention, survival curves are stretched along the age axis by a dimensionless factor.

To extract the Gompertz exponent *α* from the mortality data (Stroustrup et al. 2016), we used the corresponding survival curves and fitted them to the prediction of the Gompertz equation. The procedure is only sensitive to the behavior of the survival curves in the neighborhood of the average lifespan. This is fortunate, since a gene regulatory network’s stiffness parameter α coincides with the Gompertz exponent in this interval only. The value of the plateau mortality 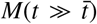 was then calculated from the tail of the cumulative hazard *m*(*t*), estimated from the raw mortality data by the well-defined Nelson-Aalen routine (Borgan 2005) with the help of the *Lifelines* package (Davidson-Pilon 2016).

Since the mortality rate reaches a plateau at late ages, the behavior of the cumulative hazard for these ages is linear and the value of 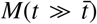 can be extracted by linear regression of the cumulative hazard on age. We calculated the cumulative hazard *m*(*t*) from the experimental data and as the prediction of the Gompertz equation and compared them in Fig. S3, where the disagreement between the two is substantial and significant, both qualitatively (exponential growth for the Gompertz equation and linear growth for the plateau mortality) and quantitatively (the cumulative hazard for the Gompertz equation is several orders of magnitude larger for late ages).

### S8 Preparation of the signature of aging for the Connectivity Map screening

To transform the list of genes associated with aging in *C. elegans* into the form appropriate for the Connectivity Map (CMAP) database (Lamb 2007; Lamb et al. 2006), we first identified human orthologs for the genes from this list using OrthoList database (Shaye et al. 2011) comprising information from four other databases: Ensembl Compara (Aken et al. 2016), InParanoid (Sonnhammer et al. 2015), NCBI HomoloGene Database, OrthoMCL (Li et al. 2003). Since, CMAP requires human genes to be presented by HG-U133A tags (Affymetrix Human Genome U133A Array), the g:Profiler database (Reimand et al. 2016) was used to map human Ensembl gene IDs to HG-U133A tags. Finally, the lists of up-and down-differentially expressed with age genes were formed and used to predict the list of prospective drugs-perturbagens using CMAP. These drugs are expected to reverse the gene-expression profiles to a younger state.

